# When to depart from a stopover site? Time since arrival matters more than current weather conditions

**DOI:** 10.1101/2020.02.05.933788

**Authors:** Sébastien Roques, Pierre-Yves Henry, Gaétan Guyot, Bruno Bargain, Emmanuelle Cam, Roger Pradel

## Abstract

On the journey to wintering sites, migratory birds usually alternate between flights and stopovers where they rest and refuel. Migration strategies are assumed to differ according to season: a time-minimization pre-breeding migration strategy towards breeding locations, and an energy-minimization post-breeding migration strategy to wintering ones. The duration of flights and stopovers determines the energy requirements and the total duration of the journey. Since migrating birds actually spend most of the time at stopovers, selection to minimize the amount of energy or time spent on migration is very likely to operate on the effectiveness of stopover rest and refueling. Here we address the relative contribution of factors to departure decisions from stopover sites during the post-breeding migration in a long-distance migratory songbird. When capture probability is low it is impossible to assess the variation in body condition over the entire duration of the stopover. To get around this, we use Time Since Arrival (TSA) as a proxy for the changes in the state of individuals during the stopover. We propose that TSA is an integrative proxy for resting, feeding and fattening efficiency. We develop a capture-recapture model to address the relationship between departure probability, estimated TSA, and weather conditions. Using a 20-year dataset from sedge warblers, we show that TSA has a larger effect on departure probability than weather conditions. Low humidity and an increase in atmospheric pressure in the days preceding departure are associated with higher departure probability, but these effects are smaller than that of TSA.

## Introduction

Each year migratory birds species commute between breeding and wintering areas (Alerstam 1990, Somveille et al. 2015). Most species cannot go from breeding to wintering grounds in a single flight of thousands of kilometers. They must stop to rest, feed and refill energy stores regularly (Alerstam 1990, Åkesson and Hedenström 2007, Schmaljohann and Eikenaar 2017, Schmaljohann 2018). The duration of migratory flights and stopovers determines the energy requirements and also the total duration of the journey. To maximize survival, individuals must optimize their journey to match with seasonal variation in food availability (and their energy requirements) at stopover sites and to arrive at wintering areas when resource availability is sufficient (Alerstam 1990, Somveille et al. 2015, 2018, 2019; Zúñiga et al. 2017, Schmaljohann 2018). Birds spend much more time at stopover sites than in migratory flight, with almost 85% of the journey spent on stopover (Hedenström and Alerstam 1997, Green et al. 2002, Schmaljohann et al. 2012, Schmaljohann 2018). Consequently, selection to minimize the amount of energy spent (energy minimization) or the total time spent on migration (time minimization) likely operates mainly on the effectiveness of stopover rest and refueling (Hedenström and Alerstam 1997, Schmaljohann 2018).

Deciding to leave the stopover site normally means a new flight of hundreds of kilometers. This movement itself, and its termination, are highly constrained by the maximal flight capacity (given the size of energy reserves and body size) and atmospheric conditions (Alerstam, 1990; Jenni & Schaub, 2003; Schmaljohann & Eikenaar, 2017; Wikelski et al., 2003). Once settled at a stopover site, the probability of departing (and the duration of the stay) depends on factors associated with the initiation of movement (i.e. to depart from the site) on fuel store, resting state, food availability, weather conditions and migratory experience (Jenni and Schaub 2003, Schmaljohann and Eikenaar 2017). Recent studies have tried to disentangle the environmental factors driving departure decision (Schaub et al. 2008, Ktitorov et al. 2010, Arizaga et al. 2011, Deppe et al. 2015, Dossman et al. 2016, Schmaljohann and Eikenaar 2017). Weather conditions are usually considered through the constraint they impose on flight. Departure decision depends on wind speed and direction (tailwind assistance reduces the cost of flight; Tsvey et al. 2007, Arizaga et al. 2011, Ma et al. 2011, Dossman et al. 2016). This decision also depends on rainfall or humidity which usually force birds to stay at the stopover site (Tsvey et al. 2007, Arizaga et al. 2011, Deppe et al. 2015, Dossman et al. 2016). Last, cloud cover influences departure decision by decreasing visibility and the ability of birds to navigate (Zehnder et al. 2001, Åkesson and Hedenström 2007).

But the internal state of the individual (resting state, fuel store, migratory experience) also largely influences departure decision: birds need to rest and reach a sufficient level of fuel store to perform the next migratory flight (Alerstam 1990, Hedenström and Alerstam 1997, Schaub et al. 2008, Goymann et al. 2010, Schmaljohann et al. 2012, Dossman et al. 2016, Schmaljohann and Eikenaar 2017, Moore et al. 2017, Anderson et al. 2019). Apart from resting to recover from extreme physical exercise and sleeping to recover from sleep deprivation during migratory flight (Schwilch et al. 2002), birds at stopover allocate most of their time and energy to foraging (Hedenström and Alerstam 1997, Cohen et al. 2014, Smith and McWilliams 2014). They are assumed to refill their energy stores as fast as possible, and to continue their migratory journey if their energy stores and weather conditions are favorable (Schmaljohann 2018). In addition, the ability to refuel and rest may depend on age or experience, which can lead individuals s migrating for the first time (juveniles) and those that already traveled to wintering areas in the past (adults) to depart from stopover sites after stays of different durations or under different weather conditions (Deppe et al. 2015, Dossman et al. 2016, Schmaljohann et al. 2018).

To our knowledge, the relative importance of individual internal state (resting state, fuel store) and weather conditions at determining departure probability has never been assessed within a single modelling framework. The different results of previous studies concerning the relationship between fuel store and departure probability could be due to different local constraints or to the migration strategy (time or energy-minimizing; Hedenström and Alerstam 1997; Schmaljohann 2018; Anderson et al. 2019). But more importantly, it could be due to the fact that fuel store can only be measured when individuals are captured (i.e., not necessarily at the very beginning and end of their stay). The lack of an effect of fuel store on departure probability could also arise because the method to jointly incorporate internal individual state and environmental covariates in the same capture-recapture (CR) framework was not available (Jenni and Schaub 2003, Schmaljohann and Eikenaar 2017). Here we define and estimate a variable that cannot be measured without using electronic devices: the time the individual has spent at the stopover site since its arrival: Time Since Arrival (Pledger et al. 2008). Our goal is to assess the influence of TSA on departure probability using a long-term dataset created when electronic devices were not available, and to include a large number of individuals and years in the analysis. Despite limitations of retrospective analyses of long-term datasets (modern electronic techniques were not available), such datasets offer interesting opportunities: (i) large sample sizes to draw inference about departure probability using statistical approaches, and (ii) long time series of observations to characterize long-term patterns and yearly deviations from deep-rooted trends. Birds need to rest after a long distant migratory flight and fattening increases with the duration of the stay at stopover (Alerstam 1990, Schwilch and Jenni 2001, Jenni and Schaub 2003, Schaub et al. 2008, Schmaljohann and Eikenaar 2017). We propose that Time Since Arrival (TSA) is an integrative proxy for the overall change in the internal state of stopovering birds. If this assumption is true, then departure probability should decrease as the TSA increases, and TSA should be a major determinant of departure probability. Importantly, TSA must be estimated using an analytical technique accounting for the daily capture probability of marked individuals (i.e., the probability of capturing an individual that is alive and present in the site). This is a methodological challenge because investigators know neither the day when a bird arrives nor the day when it departs from the site. Many capture-recapture stopover studies have relied on the assumption that birds are captured for the first and last time on the exact days of arrival and departure from the site (Schmaljohann and Eikenaar 2017). This assumption is unrealistic: the probability that a bird is captured in a given day usually lies between 0.1 and 0.2 (Schaub and Jenni 1999, Schaub et al. 2001, Moore et al. 2017), and the resulting “minimal stopover duration” (duration between first and last captures) strongly underestimates the actual stopover duration (by a factor that may be as large as 3; Schaub, Pradel, Jenni, & Lebreton, 2001). Estimating TSA requires the development of a statistical model accounting of imperfect detection probability.

Here we evaluate the contribution of estimated TSA and weather conditions to departure probability. We develop a capture-recapture model and analyze a long-term CR dataset from a long-distance migrant songbird at a stopover site. Since refueling seems to be the primary biological function of stopover (Schmaljohann and Eikenaar 2017, Schmaljohann 2018, Klinner et al. 2020), we hypothesize that TSA is be the main driver of departure probability in this long-distant migrant and that wind, humidity, cloud cover and atmospheric pressure have a secondary, but significant effect on the departure probability. Estimated TSA should be closer to the genuine duration of the individual’s stay than the time elapsed between the first and the last physical captures of the individual. If TSA integrates the overall changes in the individual internal state, this should help us detect the effect of stay duration on departure probability, if any. Moreover, we hypothesize that stopover duration differs according to migratory experience, as documented in other studies (Deppe et al. 2015) and test for differences between juveniles and adults.

## Material and methods

### Study area, sampling and dataset

The study site is the Trunvel ringing station (Tréogat, Brittany, France; 47.8964859°N, 4.3698274°W). Data from marked individuals have been collected using a standardized mist-netting protocol (including tape-luring; B. Bargain, C. Vansteenwegen, & J. Henry, 2002). Each captured bird was identified, ringed and aged. We used data collected from 1990 to 2014, from the 1^st^ to the 30^th^ of August because 90% of the captures occur in this month. The years when the number of recaptures was too low were not used in analyses (less than 10 birds recaptured only once). The Sedge Warbler *(Acrocephalus schoenobaenus)* is the most abundant species that stopovers at this site during its journey to winter quarters in sub-Saharan Africa. This 12-g songbird strictly depends on reedbeds where it essentially forages on one aphid species *(Hyalopterus prunii)* to refill energy stores (Bibby and Green 1981).

Between 1990 and 2014, 79,700 individuals have been marked (Dehorter and CRBPO 2015). Among all migrant songbirds that use this site, only a small fraction stays several days to rest and refuel (i.e. actual stopover; Warnock, 2010). The majority either continues migration by the following night, or moves to another stopover place (i.e., transients; Bächler & Schaub, 2007; Schaub et al., 2008). As we aim to study the departure probability of birds that stayed over at the site, we analyzed only capture-recapture data from birds that were caught at least twice during a season (including recaptures during the same day). Hence, the estimated stopover duration applies only to the part of the population passing by the site and that stays for at least some hours or days. The sample we used included data from 683 adults and 4927 juveniles; their latest recapture occurred at the site on average 3.4 ± 3.6 (SD) days after their first capture. The mean mass gain between first and last capture on individuals was 0.48g ± 1.48 (SD).

### Weather conditions

Weather variables expected to influence daily departure probability (between day t-1 and t) were: (i) wind (on day t-1), (iii) relative humidity (on day t-1), (iii) cloud cover during the night [i.e. between day t-1 and t; scale from 0 (no cloud) to 8 (complete sky cover)] (iv) atmospheric pressure; as birds likely perceive changes in pressure rather than pressure itself, we used the change in atmospheric pressure between day t and day t-1 as a covariate (denoted as *ΔPressure* in hPa). Depending on its direction, wind can either facilitate flight (tailwind) or increase the cost of flight (headwind). To integrate both wind effects, we computed the wind covariate as in (Arizaga et al. 2011):

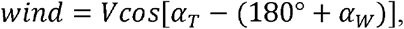

where V is wind speed (in *m,s^−1^), α_τ_* is the assumed departure direction (120° according to recovery data; note that birds don’t cross the Bay of Biscay, Dehorter and CRBPO 2015, so 120° is almost the coast direction), and *α_w_* is the direction the wind (Åkesson et al. 2002). Since birds depart on migration at the end of the day (Müller et al. 2018), we used wind speed and direction observed during a period of time starting 2 hours before sunset on day *t-1* and ending in the middle of the night on day *t*. All weather covariates were scaled prior to the analyses. The weather data were provided by the Penmarch meteorological station (47.797537, −4.374768).

### Modelling & statistical analyses

We used a formulation of the Jolly-Seber (JS) model (Jolly 1965, Seber 1965) parameterized with entry probability in the sampling area (Crosbie and Manly 1985, Schwarz and Arnason 1996). This allows modeling the arrival of birds at the stopover site and estimating stopover duration (Lyons et al. 2016, Lok et al. 2019, Roques et al. 2020). The parameters of the model are:

*φ_t_* Probability of staying in the sampling area from day *t* to *t+1*,

*η_t_* Probability of arriving at the stopover area on day *t* given that the individual was not present in the site before,

*p_t_* Probability of capturing the individual on day *t* given that the individual has arrived and has not yet left the site.

We used the Bayesian, state-space formulation of the JS model (Gimenez et al. 2007, Royle 2008). This model contains a submodel for the state process:true, partially unobservable states are “not yet arrived”, “present in the study area”, and “departed”. The model also includes a submodel for the observations (conditional on true state) directly encoded in the individual capture histories. For each individual capture history *h_i_*, the true state history is accounted for by the vector *z_i_*. This vector of binary state variables describes if an individual *i* is present in the stopover area on day *t,z_i,t_=* 1, or not, *z_i,t_* = 0.

The state process is defined as:

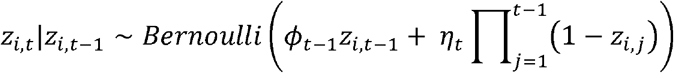

The term 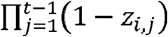 accounts for the availability of the individual to enter the stopover area and is equal to 1 when the individual has not yet entered the stopover area, and 0 when it has already entered.

As the binary observations are conditionally independent Bernoulli random variables, the link between the state and observation processes is given by the following equation:

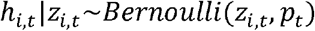

This means that if individual *i* has not yet entered the stopover area or has left *(z_i,t_*= 0), then *h_i,t_*= 0 with probability equal to 1. If *z_i,t_*= 1, then the capture history *h_i,t_* is a Bernoulli trial with probability *p_t_*, which is the probability of capturing the individual on day *t.* This formulation allows us to estimate TSA for each individual. The TSA covariate is a partially- or non-observable variable computed using the sum of true states *z_i,t_.* TSA accounts for the time individual *i* has already spent in the stopover area on day *t*:

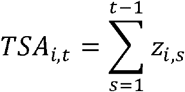

The state vector *z_i_*. also allows us to use a new formulation of the stopover duration described in Lyons et al. (2016). We computed the mean stopover duration (in days) as follows:

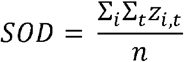

where *n* is the number of individuals and *z* the true state variable (whether individual *i* was present or not at the stopover site on day *t*).

To account for heterogeneity in detection probability among capture occasions and limit the number of parameters to estimate, we modeled detection probability as a random effect. Hence, we modeled *p* as:

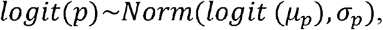

where *μ_p_*, is the mean recapture probability and σ_*p*_ the standard deviation of the random effect.

We expressed the probability of remaining at the site as a function of the previously defined weather covariates and TSA. We considered effects as ‘statistically significant’ when the estimated slope corresponding to these covariates had a 95% credible interval excluding 0 (Kéry and Schaub 2011). We analyzed the 20 years of data simultaneously, but accounted for potential differences among years (Péron et al. 2007) by means of a random year effect, where *y* is the number of the year:

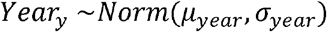

To account for the effect of the experience on an individual’s departure probability from the stopover site, we used age-dependent random effects with 2 age classes (Adult and Juvenile), where *a* is the age class:

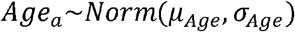

Using a logit link, the probability of staying at the stopover area between *t* — 1 and *t* was formulated as:

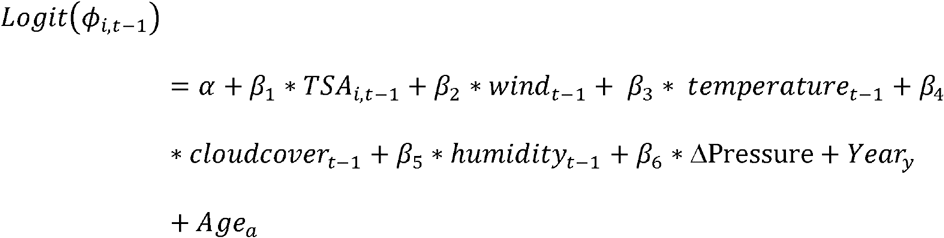

Here we included all the covariates in the model despite a non-null collinearity between most of the environmental covariates (see online Supplementary material for details). However, this collinearity appeared sufficiently small to allow the inclusion of all effects in a same model (Hair et al. 2006). Also, because TSA is computed at each occasion for each individual, TSA cannot be standardized prior to analyses. To compare the effect of TSA on departure probability to that of weather covariates, we calculated the effect of a standardized TSA by multiplying the estimated values of the TSA slopes (β_1_) by the standard deviation of all estimated TSA values.

We performed analyses with JAGS (with the package R2jags, Hornik et al. 2003; Su et al. 2015) using R version 3.6.1 (R Development Core Team 3.0.1. 2013). We used 60 000 iterations with a burnin of 30 000, and we checked chain mixing and convergence (Kéry and Schaub 2011). The JAGS code is available in the online Supplementary material.

## Results

The mean estimated stopover duration for the whole study period is 12.5 ± 2.2 days CI [12.2; 12.8], with unstructured variation among years (Figure 1). Adults (experienced birds) stay on average 1.6 days more than juveniles (naive birds) (13.8 ± 2.2 for adults, and 12.2 ± 2.1 days for juveniles; this age difference is robust through years; Figure 1). However, the 95% credible intervals and standard deviation of estimated stopover duration for juveniles and adults overlap (Figure 1) and therefore age accounts for a very limited part of the variation in stopover duration between individuals. This suggests that factors other than migratory experience and covariates accounted for in this study also determine departure probability.

**Figure 1:**
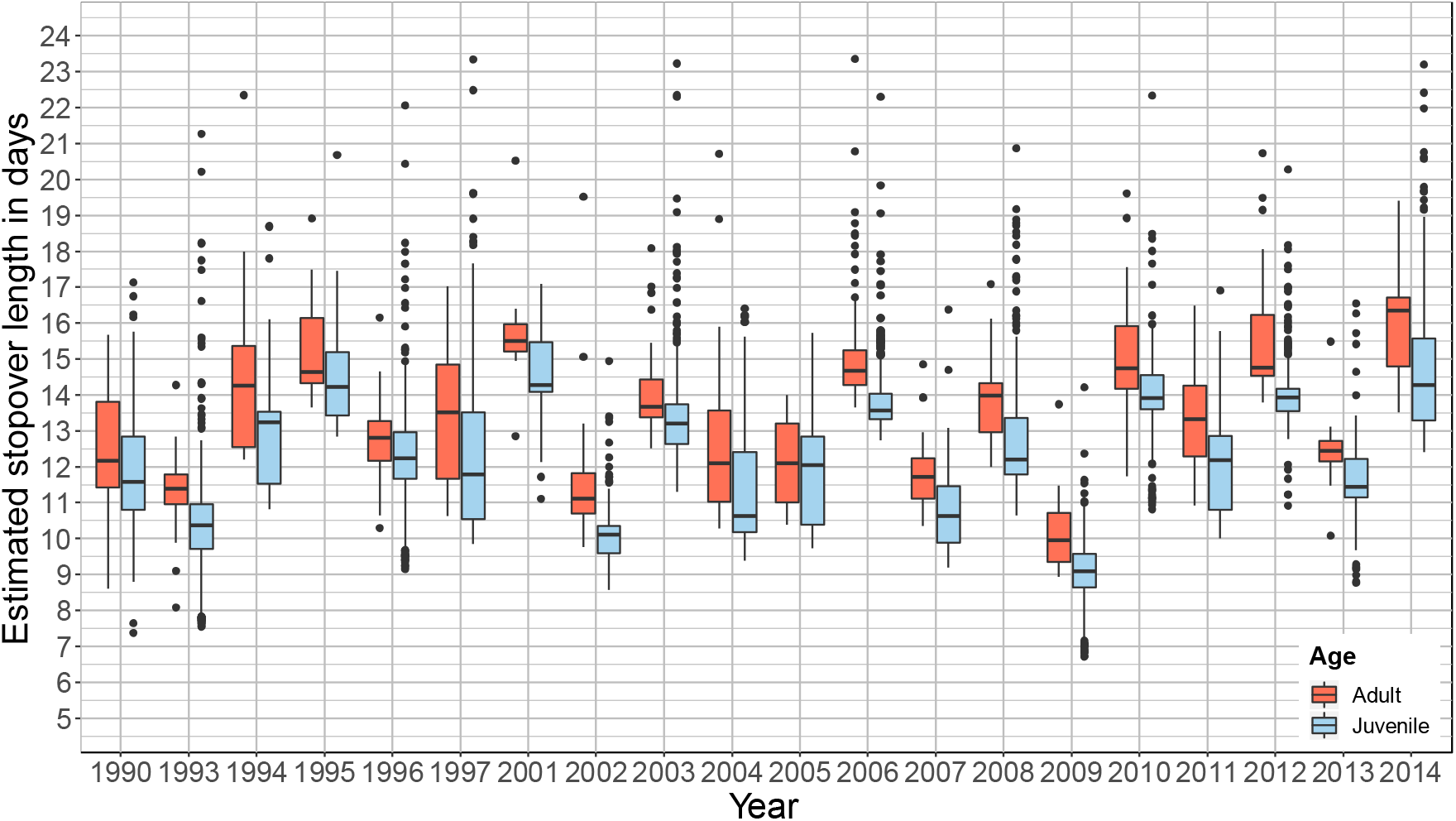
Between-year variation in individual stopover duration estimates (in days) per age category for the Sedge Warbler *(Acrocephalus schoenobaenus).*

Departure probability from the stopover site between two days (1-φ_t_, i.e. the complement of the probability of staying at the stopover site) is positively related to TSA (Figure legends and Figure 2). In other words, the longer a bird has already stayed at the site, the higher its probability of resuming migration flight by the following night. TSA is the most important predictor of departure probability compared to other variables, based on effect sizes (Table 1). TSA effect is also the effect that is estimated with the largest precision, which suggests a small variability of the TSA effect among individuals (see CI in Table 1).

**Figure 2:**
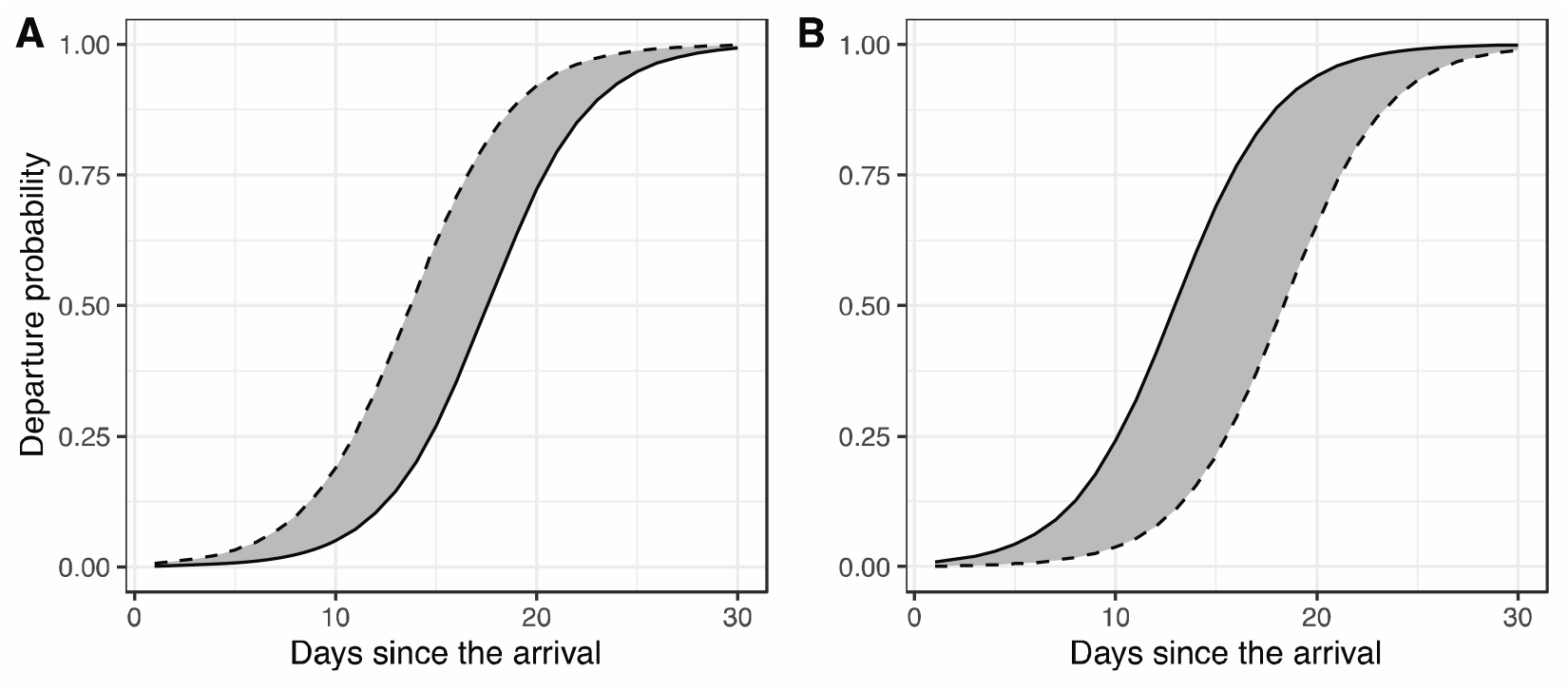
Departure probability as a function of the number of days since the bird arrived. (**A**) For different humidity conditions, dashed line: low humidity conditions (~70%); plain: high humidity conditions (~90%). **(B**) For different ΔPressure conditions, dashed line: a substantial decrease of atmospheric pressure (−5Hpa); plain: a substantial increase of atmospheric pressure (+5Hpa). The grey area represents departure probability values for humidity between 70 and 90% (a) and ΔPressure between −5Hpa and +5Hpa.

**Table 1:** Mean value, standard deviation and credible intervals (CI) for each covariate effect on the probability of staying at the stopover area. Bold: significant effects.

|  | *Mean* | *Sd* | *CI* |
| --- | --- | --- | --- |
| $(\alpha)$ *Intercept* | 6.159 | 0.175 | 5.843, 6.578 |
| $(\beta_1)$ *Time Since Arrival* | **-1.538** | **0.039** | **-1.652, -1.432** |
| $(\beta_2)$ *Wind* | 0.116 | 0.1 | -0.121, 0.196 |
| $(\beta_3)$ *Temperature* | -0.207 | 0.89 | -0.387, 0.03 |
| $(\beta_4)$ *Cloud cover* | 0.142 | 0.101 | -0.052, 0.207 |
| $(\beta_5)$ *Humidity* | **0.497** | **0.121** | **0.261, 0.735** |
| $(\beta_6)$ Δ*Pressure* | **-0.702** | **0.23** | **-1.176, -0.196** |

Both humidity and ΔPressure have an effect on the departure probability from the stopover site (Table 1). Departure probability increases with drier conditions and large changes in pressure *(Δpressure).* We did not find evidence of an effect of wind on departure probability (Table 1). We found slight evidence of a negative effect of cloud cover on the departure probability from the stopover site (Table 1), but the robustness of this result is weak because one boundary of the CI overlaps 0. Estimates of TSA and humidity effects on departure probability (Figure 2A) or of *Δpressure* (Figure 2B) show that departure probability primarily depends on TSA, and that weather covariates have a smaller influence on this probability.

## Discussion

### Effect of the Time Since Arrival on departure probability

Our statistical framework to address covariates influencing the probability of leaving a stopover site allowed us to provide evidence that TSA is the major determinant of this probability for the Sedge Warbler at our studied area. TSA is a reliable indicator of the propensity of an individual to leave the stopover site. We acknowledge that TSA is only a proxy of all the changes in the individual ‘state’*(sensu* Clark and Mangel 2000) during the stopover, but it is reasonable to think that TSA reflects the progressive change in the individual internal state. On the first day after arrival, birds are supposed to be exhausted, starving and to lack fuel stores. The longer they stay, the more opportunities they have to rest, feed and fatten (Schmaljohann and Eikenaar 2017).

TSA encompasses different functions of the stopover behavior: (i) resting after a migratory flight (McWilliams et al. 2004, Skrip et al. 2015), and (ii) reaching a sufficient level of fuel load to perform the following migratory flight (Schmaljohann and Eikenaar 2017). TSA also reflects (iii) the refueling rate, which depends on environmental conditions and physiological processes involved in refueling (Jenni and Schaub 2003, Schmaljohann and Eikenaar 2017). Here, TSA has a positive effect on departure probability: birds need to stay a sufficient number of days before leaving. For birds that do fatten, the fattening (or fuel deposition) increases through time: the longer individuals stopover, the larger their last measured body mass or fuel load (mean mass gain of 0.48g in this study; T. Alerstam, 1990; Schmaljohann & Eikenaar, 2017) and the larger the daily mass gain (Péron et al. 2007). Consequently, it is reasonable to think that birds need to rebuild a sufficient level of fuel store to perform another migratory flight (Alerstam 1990). Moreover, the resting time after a long migratory flight is apparently brief and confined to the first hours or days of the stopover (Fuchs et al. 2006, 2009; Németh 2009). This suggests that resting is not the physiological process that requires a 12-day stopover. Rather, most of the time spent in stopover is allocated to foraging in order to refill energy reserves; the latter is a long and progressive process. Hence, for the fraction of birds that stopover at this place, we believe that TSA reflects the time required by the physiological processes involved in refueling. Determining the precise relationship between TSA and fuel store will be an important area of future research in stopover ecology and more specifically for the application of the present modelling framework.

Traditional measures of fuel store such as size-scaled body mass or fat score have some limitations (extensively discussed in Schmaljohann & Eikenaar, 2017; Schwilch & Jenni, 2001), of which we want to highlight two. (i) In many long-term migration monitoring programs of marked birds, especially in old datasets, body mass was not systematically recorded at each recapture event. This drastically reduces the sample size available for long term analyses where we need a body mass measurement at each recapture. In the French dataset for the Sedge Warbler, in the twentieth century, individuals were weighed on only 60% of the capture events (Dehorter and CRBPO 2015). Nowadays (2000-2016 period), individuals are weighed on nearly all the capture events (90%). (ii) Since the probability of being captured in a given day can be low in routine trapping protocols (0.161 [0.058, 0.376] in this study; Schaub et al. 2001, Schmaljohann and Eikenaar 2017), the body mass measured at the latest capture is unlikely to be representative of the body mass that actually triggers departure. Imperfect detection probability is a common situation where modeling the individual history before the first capture is required to estimate arrival date. This imprecision in the assessment of the body mass change through time can mask the effect of body mass on departure probability in datasets where daily capture probability is low. Overall, the proposed analytical method, relying on TSA, allows analyzing individual variations in stopover duration over long time series, even in absence of biometric data, and even when body mass or fat score information are too sparse to reliably document the progress through time of the energetic state of each monitored individual.

Using TSA as a proxy for the internal state of the bird just before departure make some critical assumptions. When birds stay only few days, they may not improve condition with time spent at the stopover site because they may first continue to degrade upon arrival waiting for their digestive system to redevelop to refuel after a long-distant migratory flight (McWilliams et al. 2004). To overcome this situation, further research on the refueling process during stopover may be helpful to better model the TSA effect. The relevance of TSA may also be limited when birds relocate in the vicinity of the study site (Bächler and Schaub 2007): in this case the departure from the study site does not mean that individuals are resuming a long-distance migratory flight. Birds relocating a few kilometers away from the study site do not need to wait until their internal state improves before leaving the site. To get around these problems, birds captured only once are usually removed from capture-recapture datasets. This removes most of the transient individuals (Mills et al. 2011, Taylor et al. 2011, Sjöberg et al. 2015). Including both estimated TSA and fuel store data (body mass, fat score) into a model would be an interesting avenue to revisit previous analyses. This will help understand the contrasted results of previous studies about fuel store effects on departure probability (Tsvey et al. 2007, Schaub et al. 2008, Arizaga et al. 2011, Smith and McWilliams 2014, Schmaljohann and Eikenaar 2017). Also, other limitations, more related to modelling and data, appear when we use this type of model. It requires a large amount of recapture to test for more precise effects of the various determinants of departure probability. Interactions between TSA and date or TSA and age could have been tested. However, to correctly estimate the interaction, it is possible that it would have been necessary to drastically reduce the dataset to keep individuals captured at least three times, which does not exist for some years of our dataset. This would be interesting avenues to test on a dataset with a higher detection probability, for example with a capture-mark-resight shorebirds dataset (Lok et al. 2019).

### Effect of weather conditions on departure probability

In relatively humid days, birds tend to postpone departure. Birds wait for dryer conditions to resume their migratory flight. Humidity can be high even in absence of precipitation in Western Brittany. The negative effect of high humidity can reflect not only the inhibitory effect of rain, but also the increased flight cost when the air is very humid (Åkesson et al. 2001, Deppe et al. 2015). The probability of departing from the stopover site increases when atmospheric pressure increases between the day before the night of departure and two days before. When birds perceive an increase in pressure (indicative of improving, anticyclonic conditions), this could encourage them to resume migration flight by the following night. Unexpectedly, departure probability does not depend on wind, probably because wind was too rare and weak in the study area in August to be influential (mean wind force during the study period was 5 m.s^-1^, see Supplementary material for a summary of the weather covariates).

### Effect of migratory experience on departure probability

Juveniles (birds migrating for the first time) stay on average 1.6 days (11.6%) less than the older, more experienced birds at the stopover area. Even though this account for a limited variation in stopover duration in our study and also that there is no consensus about the age effect on stopover strategy (Hake et al. 2003, Moore et al. 2017), this result is consistent with former studies that have shown that juvenile and experienced birds behave differently regarding departure probability from stopover sites: juveniles make shorter and more frequent stopovers than adults (Reilly and Reilly 2009, McKinnon et al. 2014). However, other studies reached opposite conclusions: telemetry studies of departure decisions in songbirds in the Gulf of Mexico during autumn did not find any effect of age on the decision to cross the Gulf of Mexico, or of the selection of weather conditions (McKinnon et al. 2014, Deppe et al. 2015). Again, it is legitimate to ask whether imperfect detection probability is involved in inconsistencies among studies concerning the relationship between age and departure probability from stopover sites.

### Respective effects of TSA and weather conditions on departure probability

In this study, we show that the contribution of TSA to departure probability in a long-distant migrant is larger than that of weather conditions. This suggests that even when weather conditions are favorable to departure, birds need to stay before departing. To our knowledge, this is the first time that the contributions of these factors were estimated with a capture-recapture model. Nevertheless, our result is consistent with numerous studies that have highlighted the key role of the improvement of the individual internal state (fuel store, fuel deposition rate, body condition, body mass, resting) on departure decisions from a stopover site (see Schmaljohann and Eikenaar 2017 for a review and Anderson et al. (2019) for a recent study with telemetry tools).

The improvement of the internal state during a stopover could be indirectly related to environmental conditions. Indeed, harsh weather can decrease the ability of individuals to feed and food abundance in the stopover area. This likely leads to a decrease in the rate at which birds accumulate energy (Jenni and Schaub 2003). If this holds, weather conditions can affect TSA and stopover duration. Interestingly, here we found slight evidence of an effect of weather conditions on departure probability while TSA was taken into account in the analysis. The relationship between TSA and weather conditions during the stopover should be addressed in future work to better understand the processes involved in departure decisions from a stopover site.

Recent studies also highlighted that migration distance affects the stopover strategy of birds (Anderson et al. 2019) and that birds also behave differently between stopover sites along their journey (Schmaljohann et al. 2017). Concerning the Sedge Warbler at our studied site, we have no clues from a specific origin (controls indicate birds from Great Britain, Scandinavia, Eastern Europe; Dehorter and CRBPO 2015) or a final destination. However, as we only study one species which is strictly trans-saharian, the variability of stopover strategy induced by migration distance may be limited.

## Conclusions

We incorporated TSA, a partially hidden individual state, and weather conditions in the same capture-recapture modeling framework to disentangle the factors playing a part in the decision to depart from a stopover site. Using a long-term dataset, we showed that TSA is the main driver of departure probability (and of stopover duration) in a long-distant migrant songbird. This approach will allow investigating the determinants of stopover duration and departure probability (not only weather variables but also some hidden physiological processes accounted for by TSA) in hundreds of existing long-term datasets, where there is no, or scattered information about mass or fat score. We demonstrated the feasibility and relevance of this analytical approach using data from one site, one species and over a large period of time. Our modeling approach will have to be used with data from several species, at several sites, to assess the robustness and generality of our conclusion about the major influence of TSA on the time when individuals decide to leave stopover sites.

TSA also has broader implications outside migration ecology in situations where the time individuals spend on sites is a partially observable variable. For example, in behavioral ecology and foraging ecology, the probability of an individual changing foraging site could also depend on the time spent in a site, the number of competitors, or food availability. TSA opens large perspectives when behaviour depends on the time spent in a site, in a specific state, and when detection probability is imperfect at the time scale relevant to the research topic addressed.

## Acknowledgements

This study was made possible thanks to tens of bird ringers and hundreds of assistants who volunteered in this long-term monitoring over three decades, and the continued support of the Société d’Etude de la Protection et de la Nature de Bretagne / Bretagne Vivante. This study was supervised by the Centre de Recherches sur la Biologie des Populations d’Oiseaux (CRBPO), the Museum National d’Histoire Naturelle, the CNRS and the Ministère de l’Environnement. This work was carried out in accordance with standard animal care protocols approved by the CRBPO. Emmanuelle Cam was supported by the French Laboratory of Excellence project “TULIP” (ANR-10-LABX-41; ANR-11-IDEX-0002-02).

## Data availability statement

Data used to conduct the analyses is available on the github account of the corresponding author at the following link https://github.com/sebroques/Article_Sedge_warbler

