## Supplementary material for "When to depart from a stopover site? Time since arrival matters more than current weather conditions"

**1 CORRELATION BETWEEN WEATHER COVARIATES**

Table 1 : Pearson correlation coefficient between weather covariates (the * indicates the level of significance respectively <0.05, <0.01, <0.001)

|  | Wind | Temperature | Humidity | Cloudcover | ΔPressure |
| --- | --- | --- | --- | --- | --- |
| Wind |  | -0.245*** | 0.239*** | 0.138** | 0.41*** |
| Temperature |  |  | -0.122* | 0 | -0.07 |
| Humidity |  |  |  | 0.390*** | 0.06 |
| Cloudcover |  |  |  |  | -0.08 |
| ΔPressure |  |  |  |  |  |

**2 SUMMARY OF WEATHER COVARIATES**

Table 2 : Summary of the weather covariates during the study period

|  | Mean | Standard deviation |
| --- | --- | --- |
| Wind Speed | 5.4 m.s^-1^ | 2.6 |
| Temperature | 17.7°C | 1.65 |
| Humidity | 84.2% | 7.16 |
| Cloudcover | 3.98 | 2.75 |
| Pressure | 1015.06 Hpa | 5.52 |

**3 JAGS CODE**

Computation time on our dataset with 5610 individuals (683 adults and 4927 juveniles) and 30 capture occasions: 23 days with 60 000 iterations and 2 chains

model {

\# Priors and constraints

a~dnorm(0,0.01) #intercept

b~dnorm(0,0.01) #TSA

c~dnorm(0,0.01) #wind

e~dnorm(0,0.01)#cloudcover

f~dnorm(0,0.01)#humidity

j~dnorm(0,0.01)#Deltapressure1

celsius~dnorm(0,0.01)#Temperature

#prior on age effect

for (age in 1:2){

age_effect[age]~dnorm(0,tau_age)

}

tau_age<- 1 / (sd_age*sd_age)

sd_age~dunif(0, 2)

#prior year effect

for (a in 1:20){

year_effectdep[a]~dnorm(0,tau_dep)

}#a

tau_dep<- 1 / (sd_dep*sd_dep) #Hyperparameter

sd_dep~dunif(0, 2)

\# prior recapture probability, 1.5 relies to a mean value of recapture probability for this #dataset (previous analyses)

logit_pm~dnorm(-1.5,0.01)

tau_p<- 1 / (sd_p*sd_p)

sd_p~dunif(0, 2)

for ( a in 1:20){

for (t in 1:n.occasions) {

logit(p[a,t])<- l_p[a,t]

l_p[a,t]~dnorm(logit_pm, tau_p)

}#t

}#a

\# prior for entry probs 20=number of years

for( a in 1:20){

for (t in 1:(n.occasions)) {

eta[a,t]~dunif(0,1)

}#t

}#a

\# Likelihood

for (i in 1:M) {

\# First occasion

z[i,1] ~ dbern(eta[group[i],1])

zstop[i,1]<- z[i,1]

prod1mz[i,1]<- 1

\# Observation process

mup[i,1] <- z[i,1] * p[group[i],1] * visit[group[i],1]

y[i,1] ~ dbern(mup[i,1])

\# Subsequent occasions

for (t in 2:n.occasions) {

\# State process

zstop[i,t]<-zstop[i,t-1]+z[i,t] #update TSA

prod1mz[i,t]<-prod1mz[i,t-1]*(1-z[i,t-1])

logit(phi[i,t-1])<-a+b*zstop[i,t-1]+c*wind[group[i],t-1]+e*cloudcover[group[i],t-1]+f*humidity[group[i],t-1]+j*delta1[group[i],t-1]+celsius*temp[group[i],t-1]+age_effect[ages[i]]+year_effectdep[group[i]]

mu[i,t] <- phi[i,t-1] * z[i,t-1] + eta[group[i],t] * prod1mz[i,t]

z[i,t] ~ dbern(mu[i,t])

\# Observation process

mup[i,t] <- z[i,t] * p[group[i],t] * visit[group[i],t]

y[i,t] ~ dbern(mup[i,t])

} # t

} # i

\# stopover duration

for (i in 1:M) {

stop[i] <- sum(z[i,1:n.occasions])}

zes <- mean(stop[])

}
